# Influence of Incorporating Full-Fat Black Soldier Fly Larvae Meal (BSFLM) in Meat Type Chicken Diets on the Lower Gut Short Chain Fatty Acids profile, Gut Morphology and Intestinal Lesion Score

**DOI:** 10.1101/2024.04.04.588065

**Authors:** Jayanta Bhattacharyya, PareshNath Chatterjee, Jayanta Kumar Chatterjee, Aruna Pal

## Abstract

Though protein demand is increasing day by day but the animal protein industry on a global scale is facing the biggest challenge of replacing antibiotic growth promoters (AGP) to raise broiler chickens. Using AGP is a long-standing practice to include in poultry diets for improving the intestinal health and subsequent performance of the broiler chicken. Due to increased restrictions and bans on the usage of antibiotics, the poultry-producing community is in search of a suitable and sustainable alternative to AGPs. The present study was aimed at to evaluate and analyte the potential impacts and consequences associated with the inclusion of black soldier fly larvae meal (BSFLM) in a commercially available meat type chicken when compared to diets that either contained or lacked the presence of antibiotic growth promoters, specifically enramycin at a concentration of 8% and chlortetracycline at a concentration of 15%. Our study also assessed the influence of inclusion of BSFLM on cecal short chain fatty acids (SCFAs), and the gut health. 180 male day-old Vencobb430Y chicks (mean BW 42.52 g) (P > 0.05) were randomly allocated into isonitrogenous and isoenergetic dietary treatments in three equal groups. BSFLM group has exhibited significant higher concentration of short chain fatty acids in cecum like acetate, isobutyrate, butyrate, and total SCFAs compared to both the AGP and CONTROL groups (P<0.05). The villi height of duodenum and jejunum remained significantly higher in BSFLM supplemented birds as compared to the AGP group (P<0.05). From the analysis of frequency plots depicting the severity of intestinal lesions, it was observed that the presence of serosa and mucosa congestion in the anterior sections of the gastrointestinal tract was within the expected range for both the CONTROL and AGP groups, thus indicating that this particular physiological condition can be considered as normal in these experimental groups. In the duodenal part, incorporating BSFL meal showed significantly higher villi height and crypt depth compared with CONTROL and AGP (P<0.05). Inclusion of full-fat BSF has improved overall intestinal health and lower lesion scores compared to the CONTROL and AGP groups. The present investigation explored thepotential of incorporating full-fat black soldier fly larvae (BSFL) meal into the dietary regimens of broilers for sgnificantly bolstering the health and functionality of their gastrointestinal tract, particularly in instances where the inclusion of antimicrobial growth promoters (AGP) is deliberately omitted from said diets.

## 1. INTRODUCTION

The demand of animal protein is in increasing trend and growing at a compound annual growth rate (CAGR) of 4.92% from 2024 to 2029, from an estimated 9.41 billion USD in 2024 to 11.97 billion USD in 2029. But the animal protein industry on a global scale is encountering difficulties in satisfying the growing requirement for sustenance in an environmentally responsible manner, given the apprehensions regarding climate change, resource utilization, animal well-being, antimicrobial resistance, and the availability of safe and reasonably priced nourishment[19]. Increasing demand of animal protein has put pressure in using antibiotics for improving the growth and overall productivity of the animals. The utilization of antibiotic growth promoters in feed has been a tradition in order to improve the intestinal well-being and economic efficiency of commercial broiler chickens [29,35]. One kilogramme of chicken requires an estimated 148 mg of antimicrobials to manufacture [36]. Irrational use of antibiotic and subsequent development of antimicrobial resistance is of growing concern both in humans and animals [8,13,14,16,23,33,37,38]. Limited or no use of antibiotics in animals intended for human food has exhibited lower prevalence of antibiotic resistant bacteria in animal population. The World Health Organisation (WHO) vehemently cautions against the utilization of any form of medically significant antibiotic in animals cultivated for the purpose of consumption, including instances where such antibiotics are employed for the sole purpose of enhancing growth and averting diseases in the absence of a specific medical condition or diagnosis.

Since the imposition of a ban on the administration of antibiotics to animals intended for human consumption by the European Union in 2006, the majority of nations have either implemented prohibitions or imposed stringent limitations on the utilization of antibiotics in the rearing of these animals. Furthermore, the demand for antibiotic free animal protein has been fueled by consumers, as evidenced by the adoption of “antibiotic-free” policies by numerous prominent food chains in relation to their meat procurement.

The research is in vogue to search for an alternate growth promoter other than antibiotics. One such substitute for AGPs is feeding chickens full-fat BSFLMas a functional diet. Antimicrobial compounds, immunomodulatory substances, chitin or its derivatives, lauric acid [7], and many peptides that inhibit microbes [39,42] are reported to be present in full-fat BSFLM. Full-fat BSFLM is also a great source of fat, protein, and essential vitamins and minerals [9,15].When full-fat BSFLM was added to fully replace soybean meal (SBM) in laying hens, the caecal microbiota’s composition changed. This enabled an increase in the production of microbiota-derived short-chain fatty acids (SCFAs), like butyrate, acetate, and propionate [26]. These SCFAs have a number of positive impacts on the host in addition to their antimicrobial activity [1,30]. Even with these findings, it is not yet known if full-fat BSFLM can act as a modulator of intestinal health. When compared to a standard diet without an antibiotic growth promoter, broiler diets containing full-fat BSFLM (10–20%) have been proven to improve growth performance [11]. Higher inclusion of BSFLM does not provide an economically viable alternative to AGP in the diet of commercial broiler chickens, given the present price of BSFLM. Considering the above facts in view the present study wasdesigned to assess the impact of adding full-fat BSFL @ 1% on intestinal health, specifically on caecal SCFA level, intestinal morphology, and lesion score in a commercial broiler chicken diet by replacing commonly used antibiotic growth promoters like enramycin and chlortetracycline.

## 2. MATERIALS AND METHODS

### 2.1. Composition of BSLM

The Taiyo Group, located in Chennai, successfully raised and provided the complete-fat BSFL. All flies were cultivated in identical production conditions, utilizing the same organic waste as a substrate for their rearing. After twelve days of development, the entire larval population was collected and subjected to drying at 80 °C. A chemical analysis was performed on a sample from the same batch of BSFL to determine the values of gross energy, dry matter (DM), total ash, crude protein, crude fibre, crude fat, calcium, total phosphorus, and acid-insoluble ash. The fatty acid profile and selected minerals were assessed prior to its dietary incorporation. In order to ascertain the amino acid profile, a portion of the BSFL sample was pulverized, hydrolysed, and subsequently examined at the Evonik laboratory (Evonik Operation GmBH) using the amino acid analyser ofBiochrom Make (adopting the AOAC method for total amino acids).The dry matter (DM) was obtained by measuring the weight difference between the as such andoven dried samples(AOAC 930.15-2012). The total ash was ascertained by burning in a muffle furnace at 550 °C for four hours (AOAC 920.29-1990). To estimate total nitrogen the Kjeldahl method(AOAC 954.01-1990) was adopted, and the factor N*6.25 was used calculate crude protein (CP). Using the acid-alkali technique, crude fibre was estimated (AOAC 978.10). The crude fat percentage was determinedby ether extraction in soxhlet apparatus (AOAC 920.39). Total phosphorus was estimated using the photocolorimetric method (AOAC 965.17–1966), and calcium was estimated using the titration method (AOAC 927.02-1990).The fatty acid composition of the BSFL was determined by employing the methodology described by Clayton et al. (2012). The measurement of certain trace metals, including manganese and zinc, was carried out using ICP-OES.

### 2.2. Feed ingredients and feed composition

Apart full-fat BSFL, the feed ingredients used were subjected to proximate analysis using conventional AOAC methods: corn, soybean meal, full fat soyabean, corn gluten meal (CGM), soybean oil, and rice bran. Allix software (A-system, France) was used to design three groups of feed that were all isonitrogenous and isoenergetic, with a starter, grower, and finisher for each group. Pellet feeds (grower and finisher) and crumble (starter) were prepared using the standard manufacturing process and in order to eliminate the possibility of errors in the formulation and content of all diets, representative samples of feeds from all groups were subjected to proximate analysis once more after feeds were manufactured.

### 2.3. Birds and experimental design

One hundred and eighty male day-old Vencobb430Y chicks with a mean body weight of 42.52 g)were randomly allocated into three treatment groups using a completely randomized block design. These treatments were categorized as CONTROL, AGP, and BSFLM groups. Each group consisted of six replicate pens, each measuring 1.2mx1.2m, and was provided with a feeder, an automated drinker, and bedding made of wood shavings and paddy straw. Throughout the whole experimental trial, all the birds were raised in identical environmental conditions (lighting schedule: 23 h during the first 7 days followed by 20 h a day until harvest and a starting temperature of 32°C, which was lowered by 4 °C per week considering the age of the broilers until it reached 20 °C), and a measured quantity of feed was offered daily to each of the pens in two equal divisions. Vaccination is involved against infectious bronchitis (0 d), Newcastle disease (5 d and 20 d), and infectious bursal disease (12 d). A three-phase feeding was practiced in which the starter (1–14 d) diet was given as crumbles and the grower (15–28 d) and the finisher (29–42 d) diets were given as pellets. CONTROL feeds were made with corn, soybean meal, full-fat soybean, corn gluten meal (CGM), rice bran, and a few feed additives and supplements (Table1) and considered as basal diets. The diets of the AGP groups were identical except supplemented with 15% chlortetracycline (Zoetis) and 8% enramycin (MSD). Diets of the BSFLM group were similar to CONTROL diets except full-fat BSFL meal (Taiyo Group, Chennai) was included at 1.0% as a partial replacement for soybean meal. Full-fat BSFL meal’s chemical composition was 332.3 g/kg of ether extract (EE), 431.4 g/kg of crude protein (CP), 931.2 g/kg of diet dry matter (DM), and 5648 kcal/kg of gross energy. The nutrient digestibility, apparent metabolizable energy (AME), and apparent metabolizable energy of the full-fat BSFL meal utilized in feed formulation were previously assessed through the utilization of the INRA database.

**Table 1:**
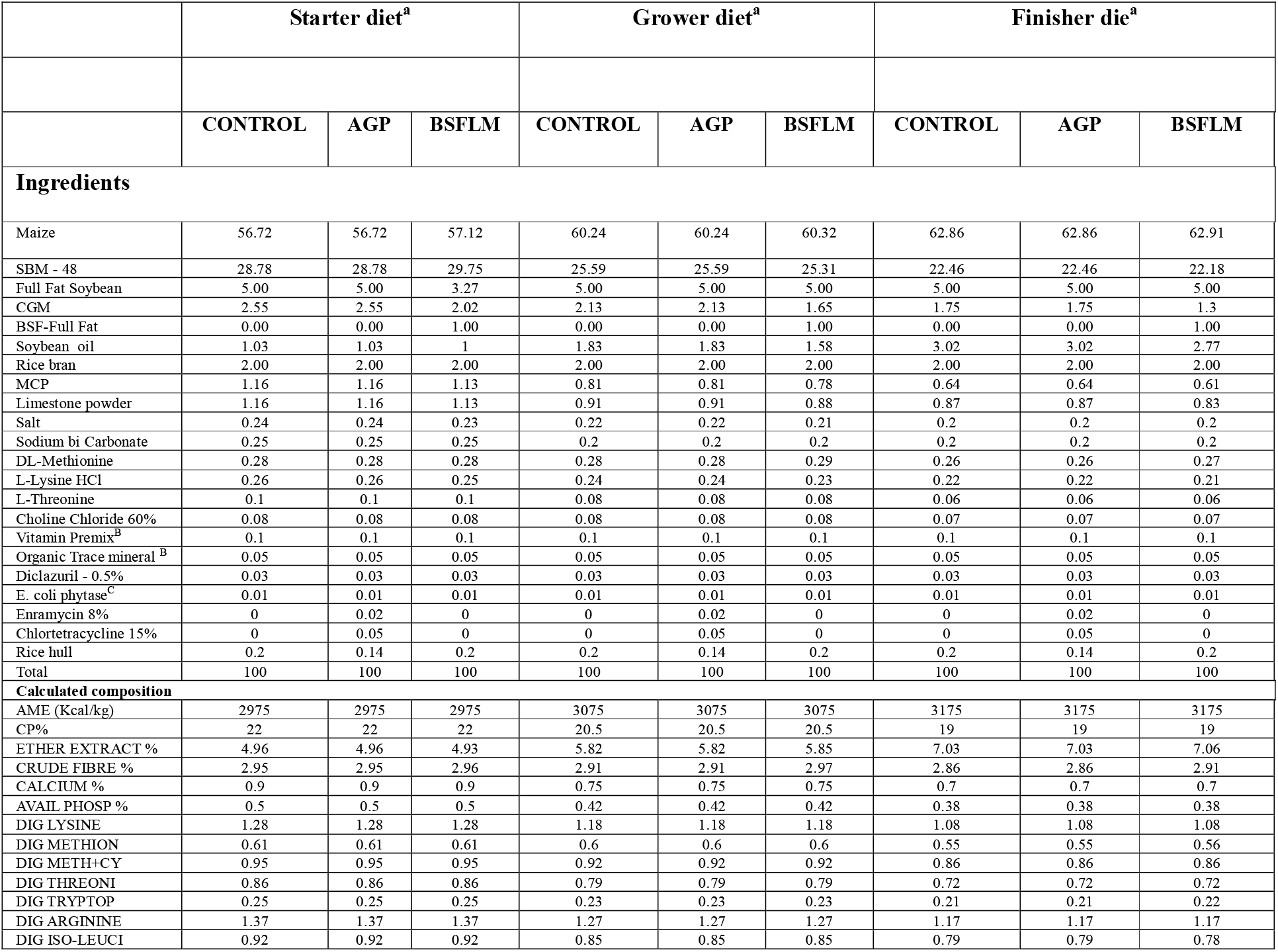
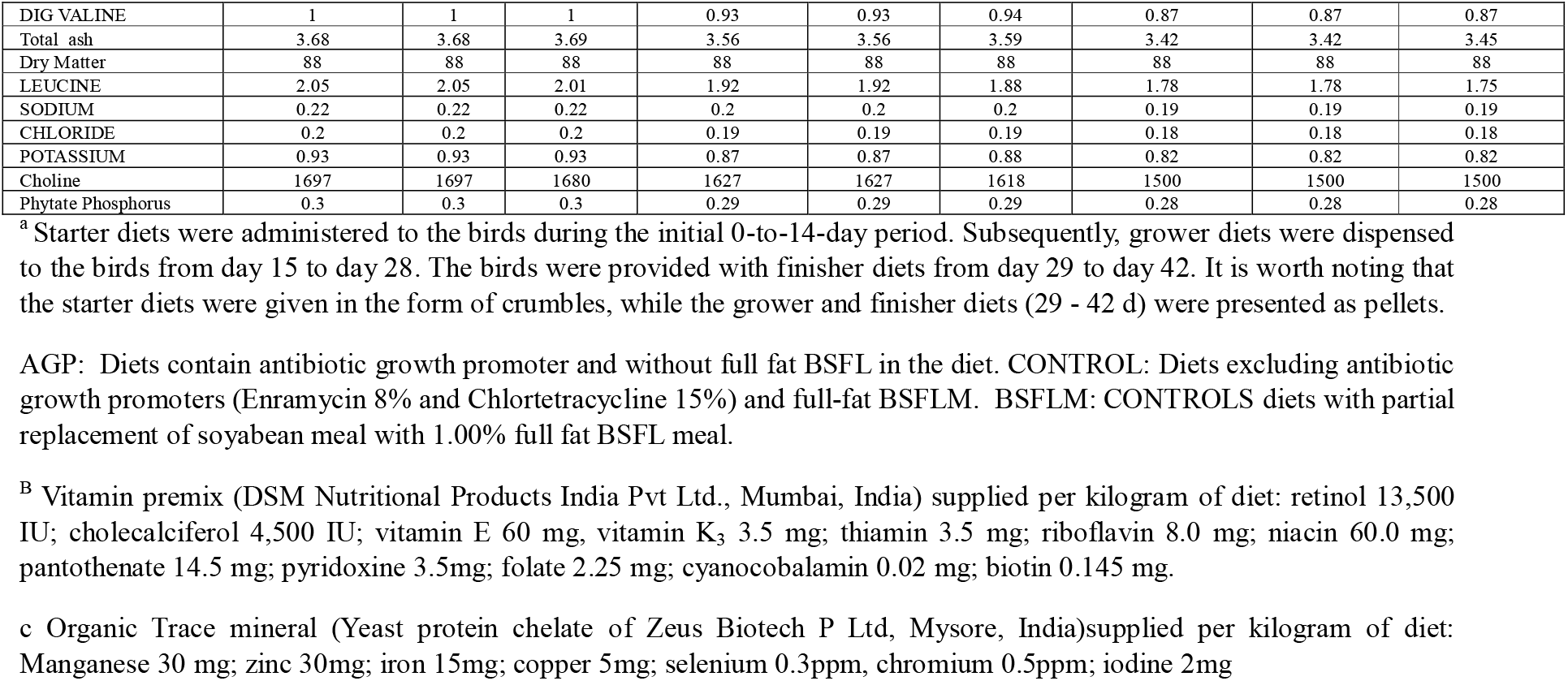
Composition and nutrients values of experimental diets.

### 2.4. Sample Collection and Analysis

On the 42^nd^ day, six birds were randomly selected per treatment (one bird from each of the replicates) and were euthanized in humane manner for sample collection by a registered veterinarian (lead author). Parts of the small intestine, which are approximately 3 to 4 cm in length, were dissected. These parts were specifically taken from the proximal duodenum, the middle of the points preceding the Meckel’s diverticulum and duodenal loop which is called as jejunum. Additionally, the mid part of the ileum was also dissected. The digesta from the ileum was taken out aseptically and preserved in phosphate buffer solution. This preserved digesta will be used for microbiota analysis, following the procedure described below. The collected samples were also preserved in 10% buffered formalin for histological analysis. As per the recommended protocol given by the manufacturer, the caecal contents of each individual bird from all three groups were immediately preserved using BioFreezeTM sampling kits (Alimetrics Diagnostics Ltd., Espoo, Finland).

### 2.4. Caecal SCFAs profiling

To calculate SCFAs, 500 mg of caecal content was collected and using pivalic acid (Sigma-Idrich, St. Louis, MO, USA) the SCFAs were analyzed as free acids in gas chromatography (GC). Caecal samples werehomogenized, centrifuged at 10,000 x g for 10 minutes at 40 °C, and then added 3 mL of ultrapure water. After homogenizing 1 mL of the supernatant with 0.2 mL of 25% metaphosphoric acid, the mixture was centrifuged at 10,000 x g for 10 minutes at 40 °C and afterwards it was placed on ice for 30 minutes. Following that, using gas chromatography (Agilent Technologies, Santa Clara, CA, USA) and a glass column filled with 80/120 carbo-pack B-DA/4% carbowax stationary phase, 400µL of the saturated oxalic acid solution and 800µL of the supernatant were examined. The helium was used as the gas and flame ionization detector.Short chain fatty acids viz., acetic acid, propionic acid, isobutyric acid, butyric acid, isovaleric acid, and valeric acid were derivatized to the respective phenyl esters by using a phenyl chloroformate reagent. The resulting esters were analyzed by Agilent GC-FID. Matrix-matched internal standard calibration with butyric-d7- and acetic-d3 acids was used in quantification.

### 2.5. Intestinal Morphology

The intestinal tissues were immersed in buffered formalin solutions with a concentration of 10% to ensure fixation. Then tissues were subjected to dehydration process in a series of alcohol solutions starting from 80% followed by 90%, 95% and finally 100%. This process lasted for a duration of 24 hours. Once dehydration was complete, the tissues were thoroughly cleaned using xylene. Finally, the tissues were embedded in paraffin blocks, which were subsequently employed for block preparation. 2 µm tissue sections were placed on glass slides and then subjected to staining with hematoxylin and eosin. Using a phase contrast microscope (Crown, Dewinter Microscope, India) with an integrated digital camera for the Dewinter microscope and DIGICAM software for image processing, the stained sections were inspected. According to Baurhoo et al. (2007), CD represents the depth of the invagination between two villi, while VH is measured from the villus tip to the villus-crypt junction. For each variable, a total of 15 measurements were taken for each sample. The average of these values was utilized for statistical analysis.

### 2.6. Small intestinal lesion score

At 42 days of age, small-intestine lesion scoring for coccidiosis and dysbacteriosis was carried out. The scoring of the small intestine lesion was conducted according to De Gussem’s (2010) method, with some modifications made for the number of birds examined.

### 2.7. Coccidia lesion scoring

Using the grading system proposed by Johnson and Reid (1970), intestinal lesions consistent with *E. maxima, E. acervulina, and E. tenella* standards were assessed for coccidia.

### 2.8. Statistical analysis

One-way ANOVA was implemented in the present study. The treatment means were compared by Tukey’s HSD test using the SPSS software package (version 21 for Windows, SPSS Inc., Chicago, IL, USA). Frequency plots were used to show the differences between the groups under intestinal lesion score analysis

## 3. RESULTS

### 3.1. Caecal SCFAs profile

Following the collection of samples, the concentration of short-chain fatty acids (SCFAs) was measured to determine the influence of feeding full-fat BSFLM on the production of SCFAs in the cecum.Propionate level didn’t differ but the concentrations of acetate, isobutyrate, butyrate, and isovalerate were significantly higher in the BSFLM group compared to other groups (P<0.05). On contrary, the AGP group had a considerably greater valerate concentration than the BSFLM group (P<0.05).

**Table 2:** Effect of partially partial replacement of SBM with BSFLM on caecal SCFA concentrations (mmol/g)

|  | CONTROL | AGP | BSFLM | Pool mean | SEM | P Value |
| --- | --- | --- | --- | --- | --- | --- |
| **SCFA** |  |  |  |  |  |  |
| Acetate | 52.36<sup>a</sup>±0.552 | 58.96<sup>b</sup>±0.742 | 64.61<sup>c</sup>±0.264 | 58.65±1.251 | 6.13 | 0.001 |
| Propionate | 6.78±0.243 | 6.43±0.147 | 6.89±0.139 | 6.70±0.110 | 0.24 | 0.201 |
| Isobutyrate | 4.83<sup>b</sup>±0.086 | 3.76<sup>a</sup>±0.315 | 7.11<sup>c</sup>±0.101 | 5.23±0.356 | 1.71 | 0.001 |
| Butyrate | 9.73<sup>a</sup>±0.150 | 10.69<sup>b</sup>±0.185 | 13.68<sup>c</sup>±0.169 | 11.368±0.418 | 2.06 | 0.001 |
| Isovalerate | 2.57<sup>a</sup>±0.113 | 3.60<sup>b</sup>±0.096 | 4.39<sup>c</sup>±0.201 | 3.51±0.197 | 0.91 | 0.001 |
| Valerate | 3.71<sup>ab</sup>±0.027 | 3.95<sup>b</sup>±0.097 | 3.32<sup>a</sup>±0.205 | 3.66±0.096 | 0.32 | 0.013 |
| Total SCFA | 79.97<sup>a</sup>±0.59 | 87.39<sup>b</sup>±0.756 | 100.00<sup>c</sup>±0.557 | 89.12±2.03 | 10.13 | 0.001 |

\*Means bearing different superscripts a, b, c differs significantly (p<0.05)

### 3.2. Intestinal Morphology

TO evaluate the impact of including the BSFLM on gut morphology, villi height (µm), crypt depth (µm), and the ratio of villi height to crypt depth (VH:CD) in the duodenum, jejunum, and ileumwere assessed using the previously described in 2.5. The histological investigation revealed adiscernible variation in jejunal morphology among the treatment groups. When compared to the AGP group, the BSFLM group’s villi remained much taller (P<0.05). However, there is no variation between the villi height and the crypt depth in the duodenum region. There are notable variations in the Jejunam region. When comparing the BSFLM group to the CONTROL group, the villi height is highest (P<0.05), although it is comparable to the AGP group (P>0.05). However, compared to the BSFLM group, the CONTROL group had the largest jejunal crypt depth (P<0.05). Ratio of villi height to crypt depth in the jejunal part in BSFLM groups showed a substantially larger value (P<0.05) than in the CONTROL and AGP groups. Surprisingly, no significant differences (P > 0.05) were observed between the groups in the iliac region. Longer villi and higher ratio of villi height to crypt depth are usually associated with higher surface area ensured for better nutritional absorption from the intestine. This suggests that feeding BSFL meals enhances intestinal morphology, which supports better gut health and ensures better feed utilization.

**Picture 1:**
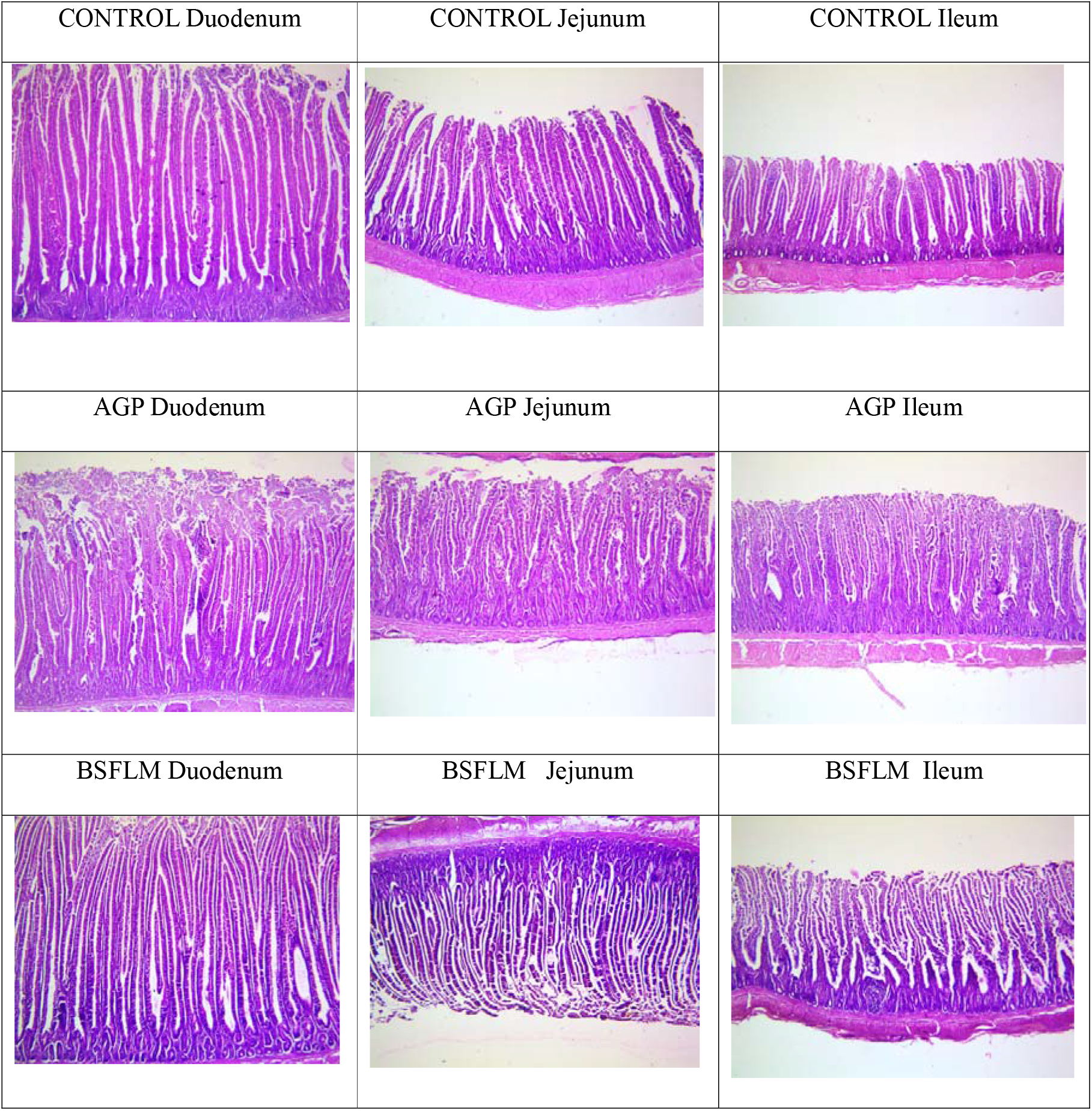
Histological images of different intestinal parts of treatment groups

**Table 3:**
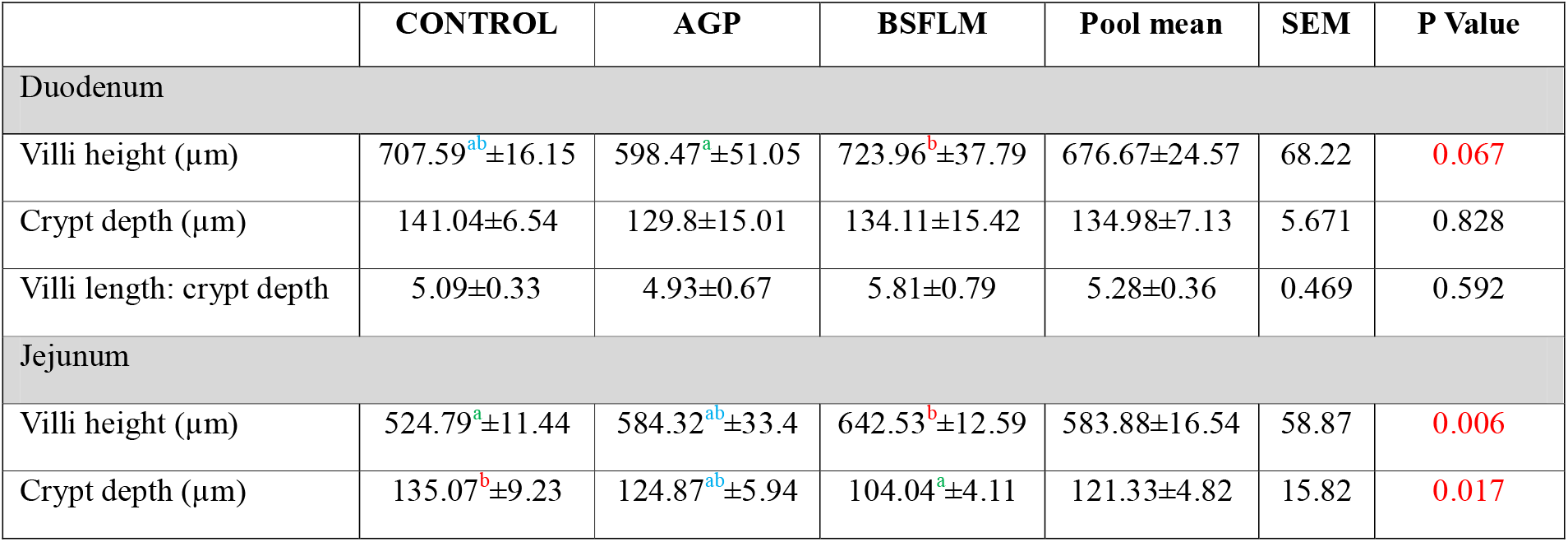
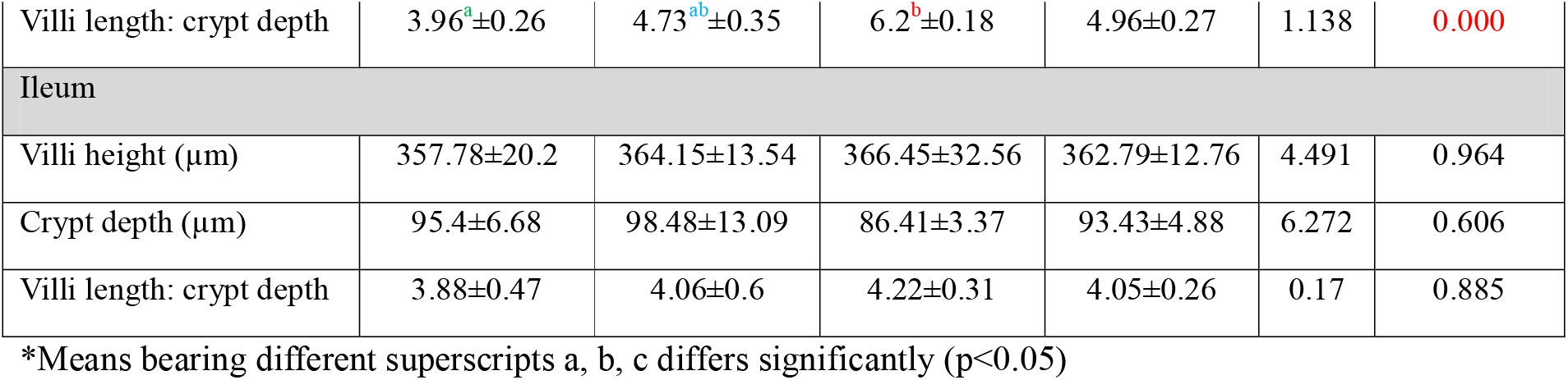
Morphometric analysis of intestinal tissues of different treatment groups.

### 3.3. Small intestinal lesion score

**Picture 2:**
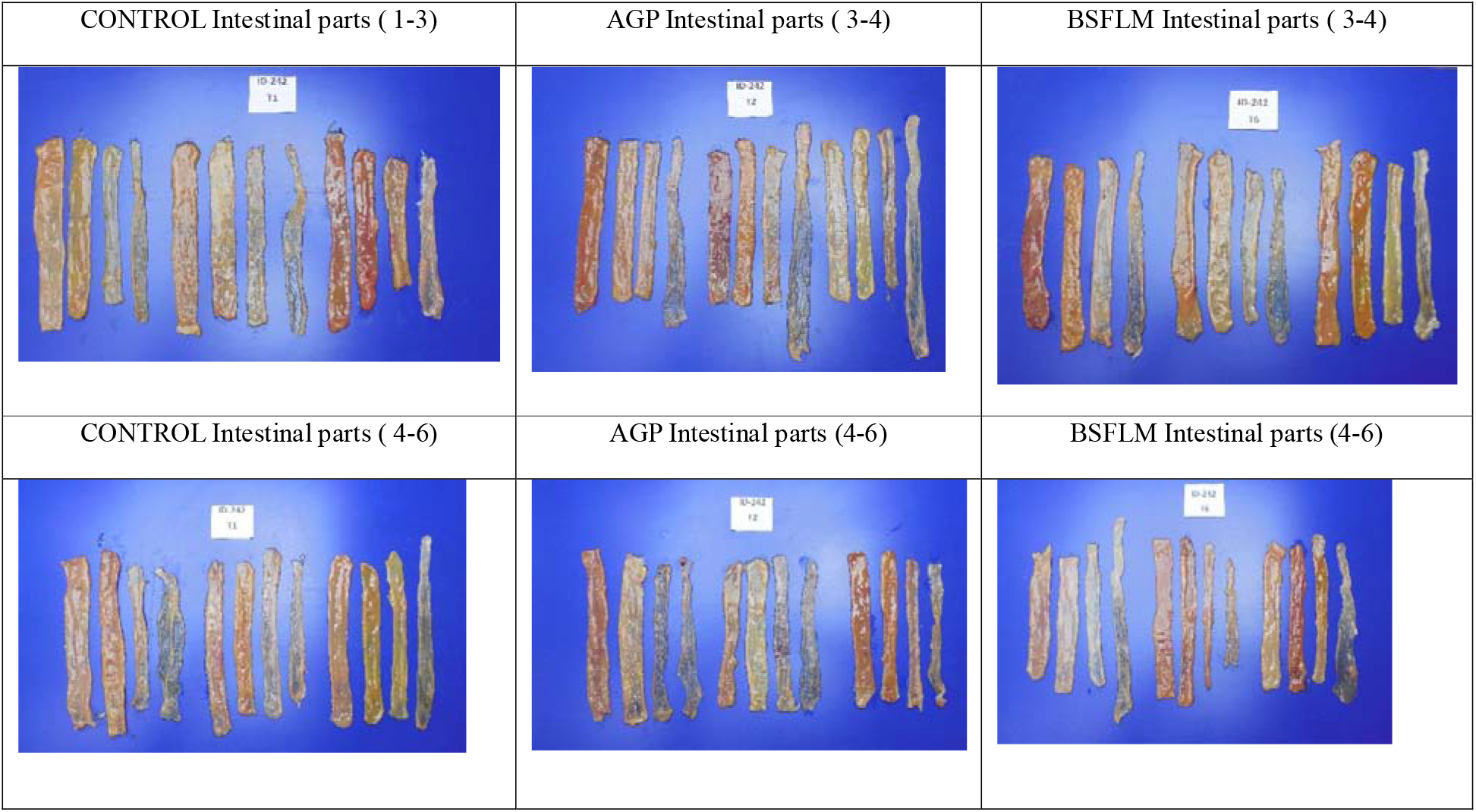
Intestinal lesion score images of different intestinal parts of experimental groups

#### 3.3.1. Cranial bacterial dysbacteriosis

According to frequency plots of the intestinal lesion score (Figs. 1.1 to 1.4), serosal/mucosal congestion of the cranial parts of the intestine was found to be nearly normal in the CONTROL and BSFLM groups, whereas on the serosal side of the gut (cranial from Meckel’s diverticulum), about 16.7% of the samples in the AGP groups had abnormally dilated blood vessels and were observed red in colour. On contrary, no variations were observedin the tonicity of the cranial sections of the intestines, which appeared to be normalbetween the groups. Compared to the control and AGP groups, 16.7% of the BSFLM samples had aberrant contents such as excessive slime, water, gas, greasy aspects, or a combination of these contents. In contrast, 16.7% of the CONTROL group’s samples revealed minor intestinal thickness or thinness in the AGP and BSFLM groups, while 33.3% of the samples showed mild thickness or thinness.

**Fig 1.1.**
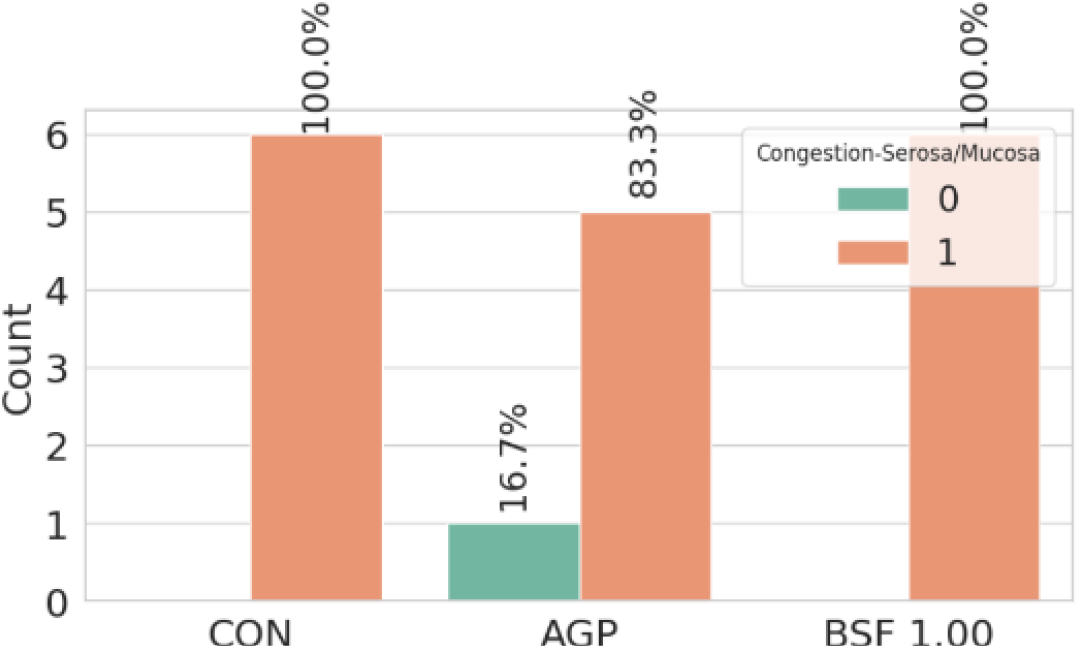
Frequency Plot of Congestion-Serosa/Mucosa for Different Treatments

**Fig 1.2.**
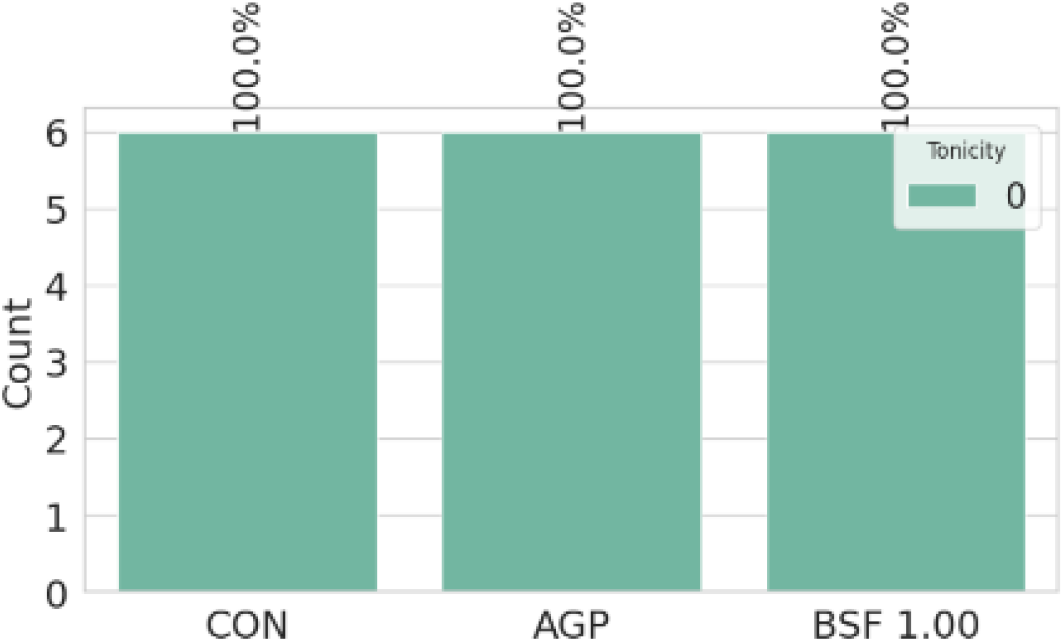
Frequency Plot of Tonicity for Different Treatments

**Fig 1.3.**
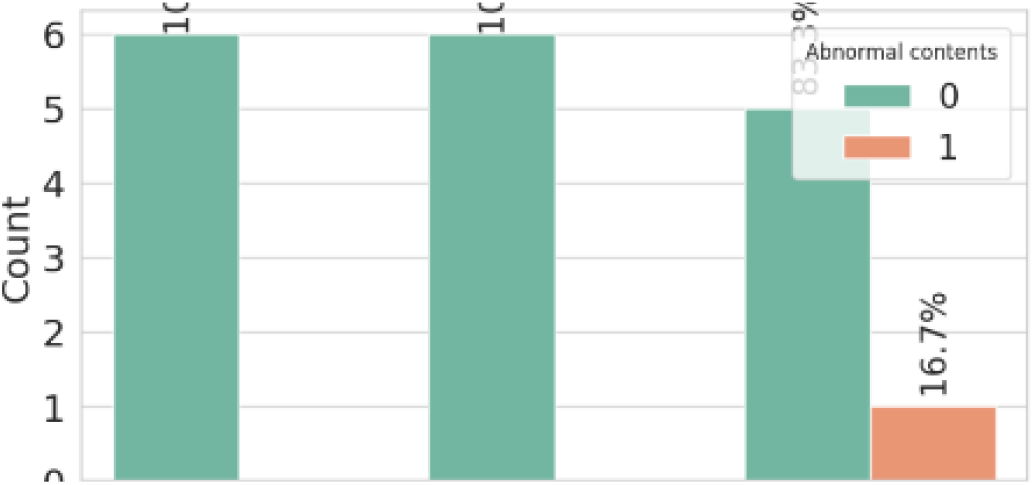
Frequency Plot of Abnormal Contents for Different Treatments

**Fig 1.4.**
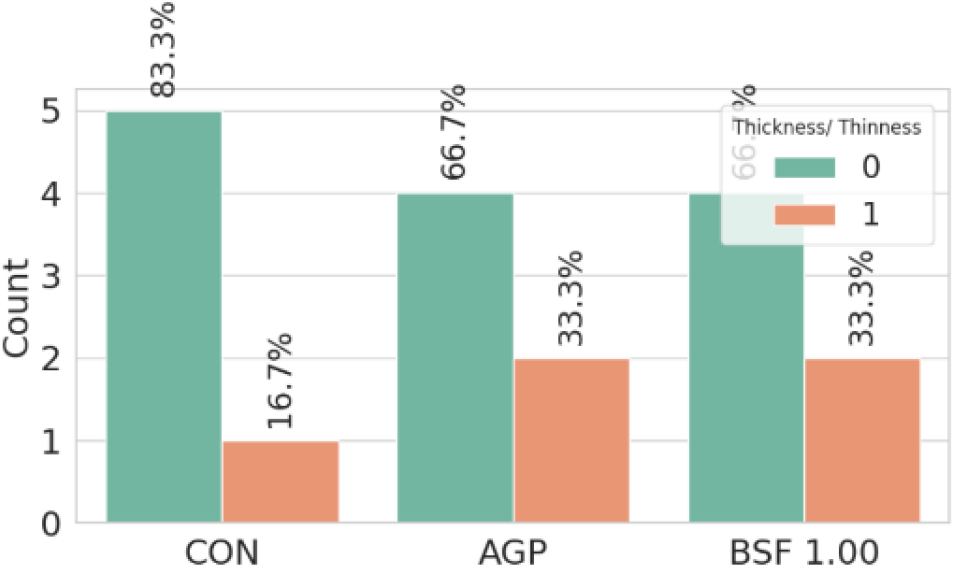
Frequency Plot of Thickness/Thinness for Different Treatments

#### 3.3.2. Caudal bacterial dysbacteriosis

From the present observations (Fig. 2.1 to 2.5) it appeared that 33.33% of the BSFLM group’s intestinal mucosa samples in the caudal region displayed abnormally dilated blood vessels and redness. In contrast, 50% of the samples from the CONTROL and AGP groups had mucosal redness and aberrantly dilated blood vessels. There was no variation in the caudal sections of the intestine’s tonicity across the groups. The CONTROL and BSFLM groups’ samples revealed aberrant intestinal contents in 33.3% of them. In contrast, AGP group which was subjected to inclusion of antibiotic growth promoters showed no aberrant intestinal contents. Within all three groups, the AGP group had a normal thickness of the intestinal tract in 33.33% of the samples. One-thirds of the CONTROL group among the groups exhibited undigested food in the colon part.

**Fig 2.1.**
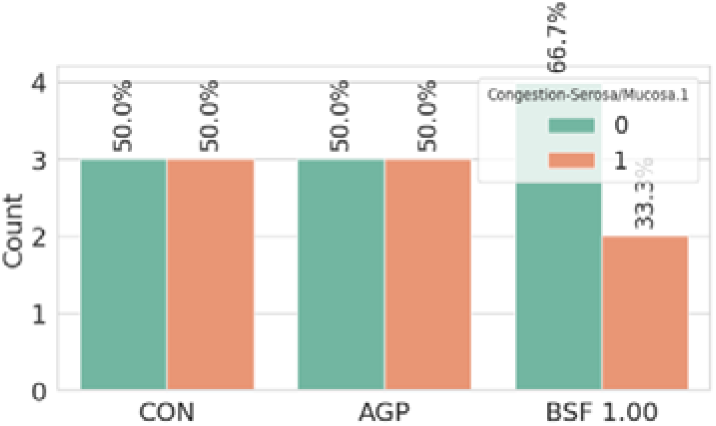
Frequency Plot of Congestion-Serosa/Mucosa for Different Treatments

**Fig 2.2.**
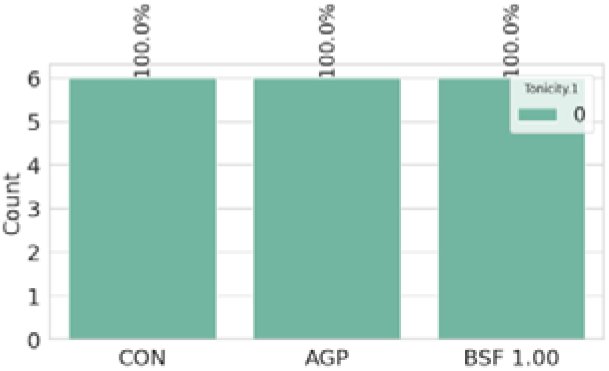
Frequency Plot of Tonicity for Different Treatments

**Fig 2.3.**
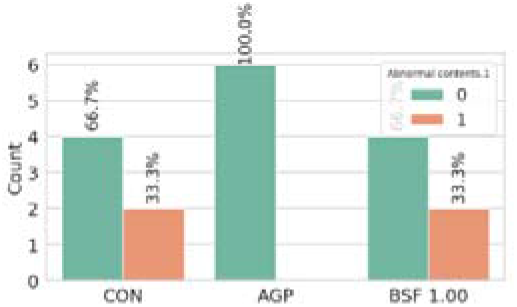
Frequency Plot of Abnormal Contents for Different Treatments

**Fig 2.4.**
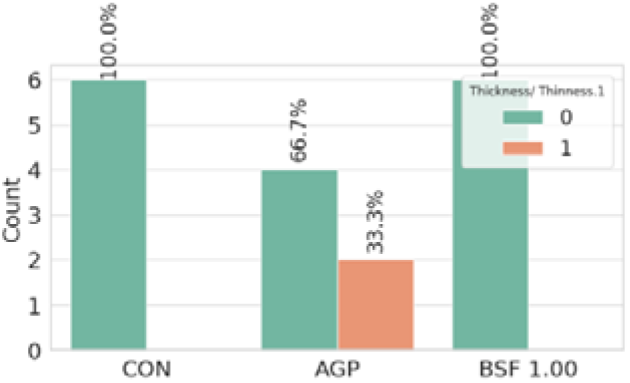
Frequency Plot of Thickness/Thinness for Different Treatments

**Fig 2.5.**
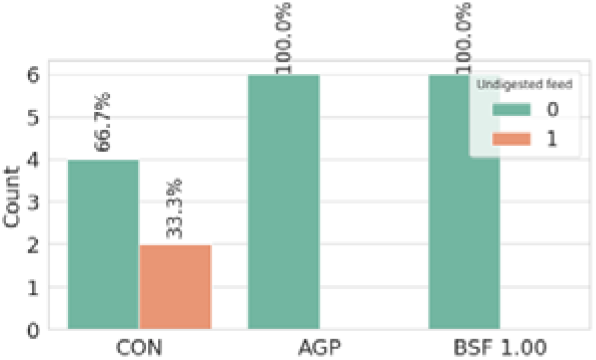
Frequency Plot of Undigested Feed for Different Treatments

#### 3.3.3. Coccidia lesion scoring

The coccidia lesion score of the samples (42^nd^ day) were presented in Fig. 1.1 to 1.5. Zero lesion scores were reported for *Eimeria acervulina*, *Eimeria maxima, Eimeria tenella*, total lesion scores, and intestinal ballooning.

**Fig 1.1.**
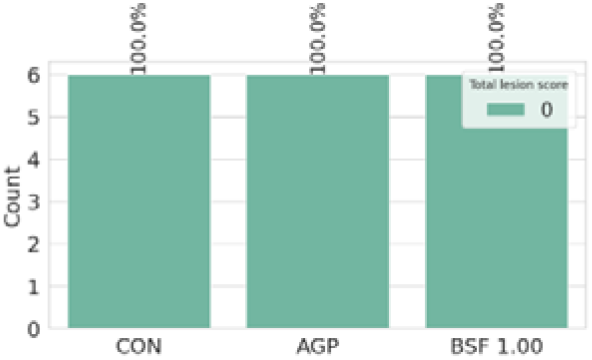
Frequency Plot of *Eimeria acervuline* for different Treatments

**Fig 1.2.**
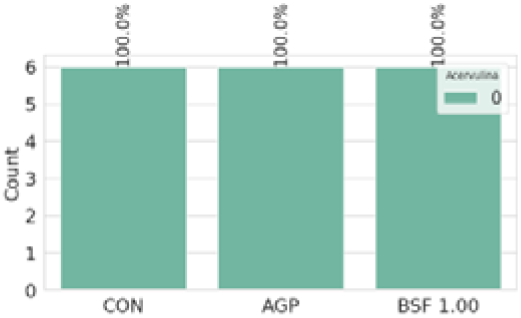
Frequency Plot of *Eimeria tenella* for different Treatments

**Fig 1.3.**
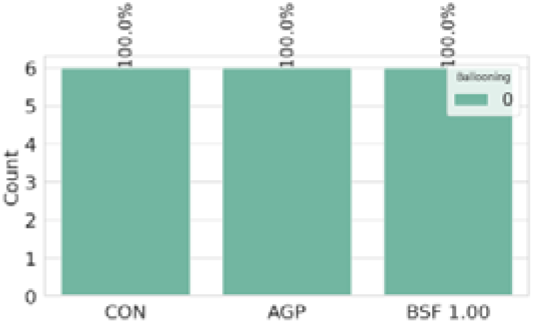
Frequency Plot of *Eimeria maxima* for different

**Fig 1.4.**
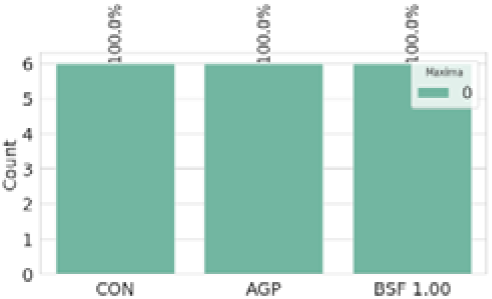
Frequency Plot of Total Lesion Score for Different Treatments

**Fig 1.5.**
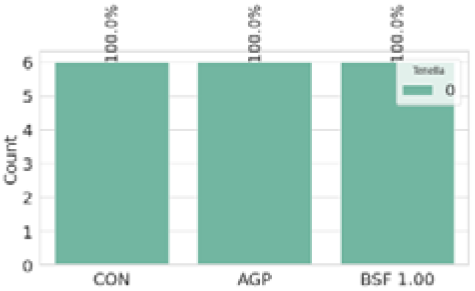
Frequency Plot of Ballooning for Different Treatments

## 4. DISCUSSION

BSFL mealpositively modulate the caecal microbiota and with concomitant rise in the height of villi when fed to broiler chickens at a rate of up to 5%. Increased inclusion levels (>15% and beyond) might be detrimental to the diversity of the existing in habitats and their beneficial metabolites [6]. Lauric acid content is greater in full-fat black soldier fly larvae meal [31]. It is experimentally found that colonization of harmful pathogens through induced infections can be effectively reduced by inclusion of specific concentration of medium chain fatty acids like lauric acid ranging from 0.1 to 1.4% [17].About 53.02% of the total fatty acids in the full-fat BSFL meal used in the experiment was lauric acid. In our study, the concentration of propionate, acetate, isobutyrate, butyrate, and isovalerate exhibited insignificantly higherpercentage in the BSFLM group over the CONTROL and AGP groups (P<0.05). Our observation is in accordance to the earlier research that showed how BSFLM, when used to replace the soybean meal (SBM), could change the composition of the cecal microbiota in a way that increased the concentration of short-chain fatty acids produced by theexisting caecal microbiota [7]. It corroborates the finding of a different study that established the abundance of microbial population residing in the caecum has a positive impact on overall health and well-being [40].

Epithelial cell renewal occurs at the depth of the crypt, and higher villi heights are linked with improved intestinal absorption of nutrients. Villi height, crypt depth, and the ratio of villi height to crypt depth are crucial in determining the absorption capacity and, in turn, the general health of the intestine.

The study showed that the BSFLM group had considerably larger villi height in the duodenum portion than the CONTROL and AGP groups (P<0.05). In contrast, the BSFLM group had the highest villi height and the ratio of villi height to crypt depth in the jejunal region of the intestine (P<0.05). On the other hand, crypt depth in the CONTROL group had noticeably higher value than in the BSFL group. There are no appreciable variations between the groups in the ileum region of the gut.

The upper small intestine, particularly the duodenum and jejunum sections, is where most nutrient absorption occurs [41]. and the intestinal health has greatly improved with the addition of 1% BSFLM as a partial replacement for SBM. This will probably have an impact on the broilers’ ability to absorb nutrients and availability of nutrients at cellular level. Researchers had anticipated that adding BSFLM, a high source of lauric acid, would promote cell renewal and, thus, improve villi length, height, and crypt depth, given that MCFA can be a direct source of energy for enterocytes [41].

Regarding the coccidial gross lesion score, no group exhibited any indications of coccidiosis. Although no group’s diet included an anticoccidial treatment, indicating that coccidiosis was well under control, it is unclear why coccidiosis was under control in the present study but might be due to maintain standard farm hygiene.

In our present study it was evident that the birds of the BSFLM group maintained a better intestinal health, both in the cranial and caudal parts of the intestine. This might be due to favourable gut microbiota modulation, which in turntriggered the production of beneficial metabolites for better intestinal health. This is most likely caused by the high amount of lauric acid in full-fat BSFLM, which inhibits*Clostridium perfringens*growth, the pathogen responsible for causing necrotic enteritis in chickens [27]. According to Benhabiles et al. (2012), chitin, the primary exoskeleton component of BSFLM, may have bacteriostatic effects and suppress Gramme-negative bacteria, including *Escherichia coli*. Furthermore, antimicrobial peptides in BSFL meal may have directly suppressed pathogenic bacteria, and chitin in BSFL meal probably served as a prebiotic to promote good bacteria like Lactobacillus and Bifidobacterium [20,34]. The altered morphology suggests that these microbiota modifications may lessen intestinal inflammation and dysfunction [25,32]. All things considered, adding BSFL seems to improve immunological homeostasis during antibiotic withdrawal, intestinal barrier integrity, and a balanced gut microbiota.

As BSFL meal is able keep broiler chicken healthy without administering antibiotic growth promoters (AGPs), it seems to be a promising approach forreducing the use of antibiotics in commercial production. In intensive broiler chicken production systems, routine antibiotic medication is frequently considered required to prevent endemic infections like as necrotic enteritis, which is associated to gut microbiota dysbiosis [3]. Producers could be able to raise broiler chicken on a commercial scale without using antibiotic growth promoter (AGP) for disease prevention or growth promotion if they use BSFL meal to improve the intestinal health.

## 5. CONCLUSION

BSFL meal seems to be one of the most promising alternative feed resources that might meet the future protein hunger of animals. The elite amino acid profile of BSFL can compete even with the soya and its précised dietary inclusion might alter the formulation cost. Feeding of BSFLproved to be beneficial for improving the lower intestinal health by increasing production of short-chain fatty acids (SCFAs). It optimized the gut health, favoured the intestinal morphology, and reduced gross intestinal lesions. The gut modulating potential of BSFL meal need to be further explored for reducing the use of antibiotic in broiler production systems. Further research is needed to refine the dietary inclusion rates and to assess benefits across diverse rearing contexts. Considering its sustainability and health advantages, BSFL meal could eventually replace a portion of conventional feeds to facilitate healthierand more responsible broiler production.

## Conflict of interest

There is no conflict of interest in this research work.

## Author’s contribution

JB, PNC: Conceptualization, design of experiment, analysis of the findings and paper drafting. JB: Wet Lab experiment execution, laboratory research, formal analysis, data recording, statistical analysis; JC, AP: Project management, paper editing, veterinary procedures as a registered Veterinary Practitioner.

## Acknowledgements

The necessary support provided by BentoliAgrinutrition India Pvt Ltd. Is gratefully acknowledged. The authors also acknowledge the support [rovided by the Agrivet Research & Advisory Pvt. Ltd for assessing some lab facility and Taiyo Group, Chennai for providing the full fat black soldier fly larvae meal.

## Notes

### Competing Interest Statement

The authors have declared no competing interest.

## REFERENCES

1. Andrew, J., Roe., Debra, McLaggan., Ian, Davidson., Conor, P., O’Byrne., Ian, R., Booth. (1998). Perturbation of Anion Balance during Inhibition of Growth of Escherichia coli by Weak Acids. Journal of Bacteriology, doi: 10.1128/JB.180.4.767-772.1998

2. Åshild, Krogdahl. (1985). Digestion and absorption of lipids in poultry. Journal of Nutrition, doi: 10.1093/JN/115.5.675

3. Batovska, D.I.; Todorova, I.T.; Tsvetkova, I.V.; Najdenski, H.M. Antibacterial study of the medium chain fatty acids and their 1-monoglycerides: Individual effects and synergistic relationships. Pol. J. Microbiol. 2009, 58, 43–47.

4. Baurhoo B., Phillip L., Ruiz-Feria C.A. Effects of Purified Lignin and Mannan Oligosaccharides on Intestinal Integrity and Microbial Populations in the Ceca and Litter of Broiler Chickens. Poultry Science. Volume 86, Issue 6, 1 June 2007, Pages 1070–1078

5. Benhabiles, M. S., R. Salah, H. Lounici, N. Drouiche, M. F. A. Goosen, and N. Mameri. 2012. Antibacterial activity of chitin, chitosan and its oligomers prepared from shrimp shell waste. Food Hydrocoll 29:48–56.

6. Biasato, I., I. Ferrocino, S. Dabbou, R. Evangelista, F. Gai, L. Gasco, L. Cocolin, M. T. Capucchio, and A. Schiavone. 2020. Black soldier fly and gut health in broiler chickens: insights into the relationship between cecal microbiota and intestinal mucin composition. J. Anim. Sci. Biotechnology. 11:11.

7. Borrelli, L., L. Coretti, L. Dipineto, F. Bovera, F. Menna, L. Chiariotti, A. Nizza, F. Lembo, and A. Fioretti. 2017. Insectbased diet, a promising nutritional source, modulates gut microbiota composition and SCFAs production in laying hens. Sci. Rep. 7:16269.

8. Carson, C., X. Z. Li, A. Agunos, D. Loest, B. Chapman, R. Finley, M. Mehrotra, L. M. Sherk, R. Gaumond, and R. Irwin. 2019. Ceftiofur-resistant Salmonella enterica serovar Heidelberg of poultry origin - a risk profile

9. Chia, S. Y., C. M. Tanga, I. M. Osuga, X. Cheseto, S. Ekesi, M. Dicke, and J. J. A. van Loon. 2020. Nutritional composition of black soldier fly larvae feeding on agro-industrial by-products. Entom. Exp. Appl. 168:472–481.

10. Clayton EH, Gulliver CE, Piltz JW, Taylor RD, Blake RJ, Meyer RG. 2012. Improved extraction of saturated fatty acids but not omega-3 fatty acids from sheep red blood cells using a one-step extraction procedure. Lipids 47(7):719–727 DOI 10.1007/s11745-012-3674-1.

11. Dabbou, S., F. Gai, I. Biasto, M. T. Capucchio, E. Biasibetti, D. Dezzutto, M. Meneguz, I. Plach_a, L. Gasco, and A. Schiavone. 2018. Black soldier fly defatted meal as a dietary protein source for broiler chickens: effects on growth performance, blood traits, gut morphology and histological features. J. Anim. Sci. Biotech. 9:49.

12. De Gussem, M. “Macroscopic scoring system for bacterialenteritis in broiler chickens and turkeys.” WVPA Meeting. Vol. 1. No. 04. 2010.

13. De Vries, S. P. W., M. Vurayai, M. Holmes, S. Gupta, M. Bateman, D. Goldfarb, D. J. Maskell, M. I. Matsheka, and A. J. Grant. 2018. Phylogenetic analyses and antimicrobial resistance profiles of Campylobacter spp. from diarrhoeal patients and chickens in Botswana. PLoS One 13:e0194481.

14. Dutil, L., R. Irwin, R. Finley, L. K. Ng, B. Avery, P. Boerlin, A. M. Bourgault, L. Cole, D. Daignault, A. Desruisseau, W. Demczuk, L. Hoang, G. B. Horsman, J. Ismail, F. Jamieson, A. Maki, A. Pacagnella, and D. R. Pillai. 2010. Ceftiofur resistance in Salmonella enterica serovar Heidelberg from chicken meat and humans, Canada. Emerg. Infect. Dis. 16:48–54.

15. Dzepe, D., O. Magatsing, H. M. Kuietche, F. Meutchieye, P. Nana, T. Tchuinkam, and R. Djouaka. 2021. Recycling organic wastes using black soldier fly and house fly larvae as broiler feed. Circ. Econ. Sustain. 1:895–906.

16. Endtz, H. P., G. J. Ruijs, B. van Klingeren, W. H. Jansen, T. van der Reyden, and R. P. Mouton. 1991. Quinolone resistance in Campylobacter isolated from man and poultry following the introduction of fluoroquinolones in veterinary medicine. J. Antimicrob. Chemother. 27:199–208.

17. Hermans, David, et al. “Application of medium-chain fatty acids in drinking water increases Campylobacter jejuni colonization threshold in broiler chicks.” Poultry Science 91.7 (2012): 1733–1738.

18. Johnson J. & Reid W.M. 1970. Anticoccidial drugs:Lesion scoring techniques in battery and floor-pen experiments with chickens. Exp. Parasitol. 28(1):30–36.

19. Judith, L., Capper. (2020). Opportunities and Challenges in Animal Protein Industry Sustainability: The Battle Between Science and Consumer Perception.. Animal Frontiers, doi: 10.1093/AF/VFAA034

20. Kabara J. J., Swieczkowski D. M., Conley A. J., and Truant J. P. Fatty acids and derivatives as antimicrobial agents *Antimicrob*. Agents Chemother. 2 1972 23–28

21. Khan, S.H. Recent advances in role of insects as alternative protein source in poultry nutrition. J. Appl. Anim. Res. 2018, 46, 1144–1157.

22. Lagat, M. K., S. Were, F. Ndwigah, V. J. Kemboi, C. Kipkoech, and C. M. Tanga. 2021. Antimicrobial activity of chemically and biologically treated chitosan prepared from black soldier fly (Hermetiaillucens) pupal shell waste. Microorganisms9:2417.

23. Liu, Y. Y., Y. Wang, T. R. Walsh, L. X. Yi, R. Zhang, J. Spencer, Y. Doi, G. Tian, B. Dong, X. Huang, L. F. Yu, D. Gu, H. Ren, X. Chen, L. Lv, D. He, H. Zhou, Z. Liang, J. H. Liu, and J. Shen. 2016. Emergence of plasmid-mediated colistin resistance mechanism MCR-1 in animals and human beings in China: a microbiological and molecular biological study. Lancet Infect. Dis. 16:161–168.

24. Luca, Borrelli., Lorena, Varriale., Ludovico, Dipineto., Antonino, Pace., Lucia, Francesca, Menna., Alessandro, Fioretti. (2021). Insect Derived Lauric Acid as Promising Alternative Strategy to Antibiotics in the Antimicrobial Resistance Scenario.. Frontiers in Microbiology, doi: 10.3389/FMICB.2021.620798

25. Martin, D.; Moran-Valero, M.I.; Vázquez, L.; Reglero, G.; Torres, C.F. Comparative in vitro intestinal digestion of 1,3-diglyceride and 1-monoglyceride rich oils and their mixtures. Food Res. Int. 2014, 64, 603–609.

26. Marzieh, Marami., A., Nobakht., Farshid, Mazlum., S., Mahdavi. (2022). Replacing soybean meal with sunflower meal in laying hens rations and its effects on cecal volatile fatty acids profile and intestinal microbial colonization. Journal of the Hellenic Veterinary Medical Society, 73(3):4373–4378. doi: 10.12681/jhvms.27060

27. Matsue, M., Y. Mori, S. Nagase, Y. Sugiyama, R. Hirano, K. Ogai, K. Ogura, S. Kurihara, and S. Okamoto. 2019. Measuring the antimicrobial activity of lauric acid against various bacteria in human gut microbiota using a new method. Cell Transplant 28:1528–1541.

28. Mauro, Antongiovanni., Arianna, Buccioni., Sara, Minieri., Ilaria, Galigani., Stefano, Rapaccini. (2010). Monobutyrine: a novel feed additive in the diet of broiler chickens. Italian Journal of Animal Science, doi: 10.4081/IJAS.2010.E69

29. Moore, P. R., and A. Evenson. 1946. Use of sulfasuxidine, streptothricin, and streptomycin in nutritional studies with the chick. J. Biol. Chem. 165:437–441.

30. Namkung H, Yu H, Gong J, Leeson S (2011) Antimicrobial activity of butyrate glycerides toward Salmonella typhimurium and Clostridium perfringens. Poultry Sci 90:2217–2222.

31. Ramos-Bueno, R.P.; Gonzalez-Fernandez, M.J.; Sanchez-Muros-Lozano, M.J.; Barroso, F.G.; Guil-Guerrero, J.L. Fatty acid profiles and cholesterol content of seven insect species assessed by several extraction systems. Eur. Food Res. Technol. 2016, 242, 1471–1477.

32. Sek, L.; Porter, C.J.H.; Kaukonen, A.M.; Charman, W.N. Evaluation of the in vitro digestion profiles of long and medium chain glycerides and the phase behaviour of their lipolytic products. J. Pharm. Pharmacol. 2002, 54, 29–41.

33. Smith, J. L., and E. D. Weinberg. 1962. Mechanisms of antibacterial action of bacitracin. Microbiol 28:559–569

34. Spranghers T, Ottoboni M, Klootwijk C, Ovyn A, Deboosere S, De Meulenaer B, Michiels J, Eeckhout M, De Clercq P, De Smet S (2017) Nutritional composition of black soldier fly (*Hermetiaillucens*) prepupae reared on different organic waste substrates. J Sci Food Agric 97:2594–2600.

35. Stokstad, E. L. R. 1950. Further observations on the “Animal Protein Factor”. Proc. Soc. Exp. Biol. Med. 73:523–528.

36. Van Boeckel, T. P., E. E. Glennon, D. Chen, M. Gilbert, T. P. Robinson, B. T. Grenfell, S. A. Levin, S. Bonhoeffer, and R. Laxminarayan. 2017. Reducing antimicrobial use in food animals. Science 357:1350–1352

37. Wang, Y., C. Xu, R. Zhang, Y. Chen, Y. Shen, F. Hu, D. Liu, J. Lu, Y. Guo, X. Xia, J. Jiang, X. Wang, Y. Fu, L. Yang, J. Wang, J. Li, C. Cai, D. Yin, J. Che, R. Fan, Y. Wang, Y. Qing, Y. Li, K. Liao, H. Chen, M. Zou, L. Liang, J. Tang, Z. Shen, S. Wang, X. Yang, C. Wu, S. Xu, T. R. Walsh, and J. Shen. 2020b. Changes in colistin resistance and mcr-1 abundance in Escherichia coli of animal and human origins following the ban of colistin-positive additives in China: an epidemiological comparative study. Lancet Infect. Dis. 20:1161–1171.

38. Ward, M. J., C. L. Gibbons, P. R. McAdam, B. A. D. v. Bunnik, E. K. Girvan, G. F. Edwards, J. R. Fitzgerald, M. E. J. Woolhouse, and C. A. Elkins. 2014. Time-scaled evolutionary analysis of the transmission and antibiotic resistance dynamics of Staphylococcus aureus

39. Xu, J., X. Luo, G. Fang, S. Zhan, J. Wu, D. Wang, and Y. Huang. 2020. Transgenic expression of antimicrobial peptides from black soldier fly enhance resistance against entomopathogenic bacteria in the silkworm, Bombyx mori. Insect Biochem. Mol. Biol. 127:103487.

40. Y., Yan., Xiaochen, Chen., Zhan, Yong, Wang. (2023). Effects of Black Soldier Fly Larvae (Hermetiaillucens Larvae) Meal on the Production Performance and Cecal Microbiota of Hens. Veterinary sciences, 10(5):364–364. doi: 0.3390/vetsci10050364

41. Zentek J, Buchheit-Renko S, Ferrara F, Vahjen W, Van Kessel AG, Pieper R. Nutritional and physiological role of medium-chain triglycerides and medium-chain fatty acids in piglets. Animal Health Research Reviews. 2011;12(1):83–93. doi:10.1017/S1466252311000089

42. Zhang, S., P. Xiong, Y. Ma, N. Jin, S. Sun, X. Dong, X. Li, J. Xu, H. Zhou, and W. Xu. 2022. Transformation of food waste to source of antimicrobial proteins by black soldier fly larvae for Défense against marine Vibrio parahaemolyticus. Sci. Total Environ. 826:154163.

